# Parasite-microbiota interactions potentially affect intestinal communities in wild mammals

**DOI:** 10.1101/076059

**Authors:** Tuomas Aivelo, Anna Norberg

## Abstract

Detecting interaction between species is notoriously difficult, and disentangling species associations in host-related gut communities is especially challenging. Nevertheless, due to contemporary methods, including metabarcoding and 16S sequencing, collecting observational data on community composition has become easier and much more common. We studied the previously collected data sets of intestinal microbiota and parasite compositions within longitudinally followed mouse lemurs by analysing the potential interactions with diversity metrics and novel joint species distribution modelling. Both methods showed consistent statistical association between certain parasite species and microbiotal composition. Both unicellular *Eimeria* sp. and cestode *Hymenolepis diminuta* had an effect on diversity of gut microbiota. These parasite species also had negative associations with several bacterial orders. In comparison, closely related species *H. nana* did not have an effect on diversity, and it had positive associations with several bacterial orders. Our results reveal potential interactions between some, but not all, intestinal parasites and gut microbiota. While environmental variables explained almost half of the total variation, of which almost half could be explained by traits of parasites and microbiota, there were no clear patterns regarding mouse lemur individual variables explaining variation in the occurrence patterns of parasite and microbiota significantly. Our results provide new hypothesis for interactions between and among parasites and microbiota to be tested further with experimental studies.

## Introduction

Interaction between species is one of the key determinants of the spatial and temporal dynamics of species communities (Ings *et al.* 2009). Communities within host individuals are no exception: the multitude of interactions between intestinal organisms, both beneficial and detrimental to their hosts, affect both ecology and evolution of these symbiont communities (Petney & Andrews 1998; Pedersen & Fenton 2007; Rigaud, Perrot-Minnot & Brown 2010; Glendinning *et al.* 2014).

Microbiota normally exceed macroscopic parasites in number, species diversity and biomass. Thus, it is not only plausible, but probable, that microbiota interacts with intestinal parasites in many ways, affecting both invasions of new parasite species and their ability to colonize the intestine and within-host dynamics of parasites (Hayes *et al.* 2010; Berrilli *et al.* 2012). Indeed, e.g., a nematode parasite of mice, *Trichuris muris*, requires microbiota interactions for successful establishment in host intestine (Hayes et al. 2010).

Interactions between different parasite species have been studied extensively in laboratory experiments (Graham 2008; Knowles 2011), but it is notoriously difficult to identify between-species interactions in observational studies (Fenton, Viney & Lello 2010; Fenton *et al.* 2014). While many studies have found random parasite assemblages indicating no interactions (Poulin 1996; Behnke 2008), some studies have also found interactions between parasites, both positive and negative (Lello *et al.* 2004). One successful way of combining an experimental approach to free-living host communities is community perturbation experiments, which have revealed parasite interactions (Knowles *et al.* 2013; Pedersen & Fenton 2015).

The experimental study of interactions between microbiota and parasites has become more feasible in recent years (Reynolds, Finlay & Maizels 2015). There has also been some observational studies both in humans and in wildlife which have looked into the interaction between microbiota and parasites — some studies have not found any microbiota changes on infection (Cooper *et al.* 2013; Baxter *et al.* 2015), while others did (Lee *et al.* 2014; Kreisinger *et al.* 2015; Baxter *et al.* 2015; Maurice *et al.* 2015). In all of the cases with significant changes, infections have been linked to higher diversity of microbiota, but the effects on bacterial OTUs have been parasite species specific.

We studied parasite and microbiota occurrence in free-living mammals longitudinally, acquiring several samples from same individuals to increase the reliability of the sampling. While both intestinal bacterial and parasite communities have been studied for a long time, new sequencing technologies have brought unprecedented resolution and coverage to identifying intestinal community composition. Nevertheless, intestinal communities have not been studied as a whole, but rather in different taxonomical groups. These groups relate to different research methods used to identify viriome (Breitbart *et al.* 2003), microbiota (Eckburg *et al.* 2005), unicellular eukaryotes (Bass *et al.* 2015) or helminth communities (Tanaka *et al.* 2014; Aivelo *et al.* 2015). Our work combines two separate metabarcoding approaches, nematodes and microbiota, and supplements it with morphologically identified parasites: other helminths (cestodes), unicellular eukaryotics *(Eimeria)* and ectoparasites (lice and ticks).

Many community ecological questions, such as the strengths and directions of species interactions, require the joint analysis of organismal abundances, and if these organisms are identified using modern tools like metabarcoding, the number of taxa in a community can be in the thousands. Nevertheless, it is possible to fully specify joint statistical models by using multivariate extensions of generalized linear mixed models (Warton *et al.* 2015). Modern joint species distribution modelling approaches allow the study of association patterns between species, while also studying their environmental responses, and thus teasing the two apart. Latent variable models are an especially exciting tool that has recently been applied for ordination as well as studying the factors driving co-occurrence (Warton *et al.* 2015). In our study, we use a Bayesian hierarchical generalized linear modelling framework to analyse species environmental responses (Ovaskainen *et al.* 2016a; b). Our models include latent variables, which model the residual co-occurrence patterns in our focal communities, quantifying hypothetical species association patterns, as well as specific parameters modelling the effects of species traits on the species responses to their environment, accounting also for phylogenetic relationships between the species (Abrego, Norberg & Ovaskainen 2016b).

Our aim was to explore associations within parasite species and between parasites and microbiota by mark-recapturing the wild rufous mouse lemur *(Microcebus rufus)* population and collecting fecal samples and related metadata. Our specific research questions were: 1) are parasites associated with each other, 2) are different parasite species associated with different microbiota compositions, and 3) which, if any, host variables affect the parasite community and microbiota composition.

## Materials and methods

### Sample collection

We followed rufous mouse lemur *(Microcebus rufus)* population in Ranomafana National Park in southeastern Madagascar (21°16’ S latitude and 47°20’ E longitude). The national park consist of lowland to montane rainforest between 500 and 1500 meters of elevation. We collected samples and data for nematodes, cestodes, eimeriids and ectoparasites from September to December 2011 and 2012 and microbiota from September to December 2012 along previously described protocol (Aivelo *et al.* 2015; Aivelo, Laakkonen & Jernvall 2016) Shortly, we trapped mouse lemurs nightly on two transects, the first within the park boundaries in secondary forest and the second in peripheral zone in highly degraded campsite area. We measured mouse lemurs, collected samples and tagged previously unseen mouse lemurs for later identification with microchips. Animal handling procedures were approved by the trilateral commission (CAFF/CORE) in Madagascar (permits: 115/10/MEF/SG/DGF/DCB.SAP/SCBSE,203/11/MEF/SG/DGF/DCB.SAP/SCBSE and 203/12/MEF/SG/DGF/DCB.SAP/SCBSE) and the research ethics committee at Viikki campus, University of Helsinki.

We quantified the number of ectoparasites on mouse lemur ears. The fecal sample was divided into four parts: the first was used to identify nematodes, the second for microbiota analysis, the third for eimeriid quantification and the fourth for cestodes. We placed the cestode sample on flotation liquid (saturated MgSO4 solution, c. 0.38 kg/l) and used McMaster’s chamber for the quantification. We took photos on cestode eggs and identified the morphospecies. We placed eimeriids in open vials in 2.5% potassium dichromate solution to allow their sporulation, moved to flotation liquid 10 days later and quantified them. We took photos of the eimeriids for identification of morphospecies. The collection and analysis of nematode and microbiota data has been previously described in detail (Aivelo et al. 2015, 2016, respectively). Shortly, we isolated nematodes with Baermann’s method, isolated their DNA and amplified a part of their 18S gene. These amplicons were sequenced with 454 sequencing (454 Life Sciences, Bradford, CT, USA), grouped in operational taxonomic units with Séance pipeline (Medlar, Aivelo & Löytynoja 2014) and processed into putative parasite species. For microbiota, we amplified V1-V2 region of 16S gene, sequenced it with MiSeq (Illumina Inc., San Diego, CA, USA) and used mothur MiSeq SOP (Kozich *et al.* 2013) to identify the OTUs.

### Microbiota diversity

We analysed the effects of different parasite species on both microbiota alpha diversity (as measured by inverse Simpson index) and richness (number of OTUs) by modelling these variables with different parasites species, host sex, trapping site, host age (divided into three age classes by protocol set by Zohdy et al. 2014), host condition (as in Rafalinirina et al. 2015), host aggressiveness, amplification batch, sequencing batch and sampling week as being explanatory variables. We used linear mixed models with the R package ‘nlme’ (Pinheiro *et al.* 2013) with mouse lemur individual as random variable and started with the complete model. If the sequencing or amplification batches did not have a significant effect, we dropped them from the model as they are strongly correlated with sampling week. We also dropped non-significant variables one-by-one from the model to see if our models were robust. This did not affect which variables were statistically significant. We also explored the effects of parasites on beta diversity and included this analysis in Appendix 2.

### Joint species distribution modelling with latent variables

We fitted a statistical joint species distribution model, combining information on environmental covariates, species traits and phylogenetic constraints, as well as the sampling study design. We fitted four models in total. Using only the parasite data for years 2011 and 2012, we fitted *i)* a model constrained with environmental covariates, species phylogenies and traits, and the sampling design included as latent variables, and *ii)* an unconstrained model, with only sampling unit level latent variable. Using the combined parasite and microbiota data for only 2012, we fitted *iii)* a model constrained with environmental covariates, species phylogenies and traits, and the sampling design as latent variables, and *iv)* an unconstrained model, with only sampling unit level latent variable. In all cases, we modelled the response community data matrix using the Bernoulli distribution and the probit link function. We fitted all the models with Bayesian inference, using the posterior sampling scheme of Abrego *et al.* (2016b). More details and applications of the modelling framework used can be found also in (Abrego *et al.* 2016a; Ovaskainen *et al.* 2016a; b). We provide the the full description of the model, including assessment of model fit, in Appendix 3, as well as the prior distributions assumed in the Bayesian analysis. Below we describe the variables used in the different models.

### Parasites

For models *i)* and *ii*), we used the presences and absences of the parasites found in the mouse lemurs during years 2011-2012 as the response matrix. For model *i)*, as environmental covariates we included the sex, age, aggressiveness and general condition of the lemurs, and with males we also accounted for the size of their testis (and considered females as individuals with extremely small testis size). We also included the time of sampling (week) and its quadratic form (week^2^) to account for the effect of seasonality. As species traits, we included whether the parasite has a direct or non-direct life cycle and whether it is an endo- or ectoparasite. In order to account for possible phylogenetic correlations in the species responses to their environment, we included species phylogenetic constraints in the model (for details, see Abrego *et al.* (2016b) and Appendix 3). We constructed the phylogenetic relationships from the taxonomic tree with five levels: domain, kingdom, superphylum, phylum and species, assuming equal branch lengths. Finally, we included random effects, which also model the co-occurrence among species, at the levels of individual lemurs, transects and year of sampling, using a latent factor approach (Abrego *et al.* 2016b; Ovaskainen *et al.* 2016a).

### Microbiota and parasites combined

For models *iii)* and *iv)* we used the presences and absences of both parasites and microbiota found in the lemurs in year 2012 as the response matrix. To avoid overrepresentation of very rare OTUs, we considered only OTUs with >9 amplicons as presences. Then to avoid sequencing and OTU picking errors, we considered the OTUs present, if there were in total >99 amplicons in at least two lemur individuals. After this, we constructed the final response matrix as presence and absence at the level of orders. For model *iii)*, as environmental covariates we included the same characteristics of the lemurs as with the parasite model *i)*, and in addition, we included whether the taxon is a parasite or part of the microbiota and microbiota was considered as having neither direct nor indirect life cycle. We constructed the phylogenetic relationships with five levels assuming equal branch lengths: domain, kingdom, phylum, class and order. Since the occurrences were modelled at the level of orders for the microbiota, but at the level of species for the parasites, we set the phylogenetic distance between the two hymenolepidid species in the phylogenetic correlation matrix to 0.99. We included latent random effects at the levels of individual lemurs and transects.

### Unconstrained models

As a point of comparison, for both data sets, we fitted unconstrained models *ii)* and *iv*), where we only included a sampling unit random effect, which models the variation in species occurrences and co-occurrences at the level of individual samples, obtained from individual lemurs, and no environmental covariates, phylogenetic constrains, nor traits. Thus, the variance across sampling units in the species responses is explained with the latent variables. By comparing the results for the constrained and unconstrained models, we can separate the associations that are solely due to the (dis)similar habitat requirements (e.g. when two species share the same habitat preferences, and hence co-occur more often than expected by random) or hidden by the (dis)similar habitat requirements (e.g. when two species share the same habitat preferences, but even after accounting for this, they still co-occur more often than expected by random) from the associations immune to the effects of the explanatory variables (i.e. we see the same association patterns regardless of the inclusion of the explanatory variables). This approach is analogous to comparing a constrained and an unconstrained ordination, with the difference of our approach being model-based (see e.g. Hui et al. 2015, Warton et al. 2015).

### Variance partitioning

Variation partitioning provides means to assess the explanatory power of different explanatory variables in relation to the same response variables, and hence give insight to which environmental variables are the most influential ones (Borcard, Legendre & Drapeau 1992). For the constrained models *i)* and *iii)*, we partitioned the variation explained by the model into the part explained with fixed effects and random effects. Moreover, we separated among the fixed effects the variation explained with covariates related to lemurs and to seasonality, as well as the share of variation explained by the traits. We also differentiated between the variation explained at different levels of random effects.

## Results

We collected complete parasite and metadata for 281 samples in two years, 2011 and 2012, and combined parasite and microbiota data for 80 samples from 2012. Prevalences for different parasites varied from 1 to 72% (Table 1). All observed lice were *Lemurpediculus verruculosus*, and all ticks belonged to *Haemaphysalis lemuris.* We identified cestodes based on shape of eggs to two distinct species *Hymenolepis diminuta* and *H. nana.* Two genotyped adult specimens also validated the identification of cestode species (Voitto Haukisalmi, pers. comm.). Eimeriids belonged to one morphospecies and nematode putative species were grouped as in Aivelo et al. (2015).

**Table 1:** Prevalence of different parasites in two years, including prevalence and absolute numbers of infected mouse lemur samples. The names of putative nematode species are based on Aivelo et al. (2015). In 2011 we had a total of 100 samples from 44 individuals, while in 2012 we had 181 samples from 57 individuals.

|  |  | 2011 |  | 2012 |  |
| --- | --- | --- | --- | --- | --- |
|  |  | Prevalence (%) | n | Prevalence (%) | n |
| Nematodes | PS1 (“*Strongyloides*”) | 34 | 34 | 72 | 131 |
|  | PS2 (“*Caenorhabditis*”) | 14 | 14 | 42 | 76 |
|  | PS3 (“Strongylida”) | 2 | 2 | 1 | 2 |
|  | PS4 (“Chromadorea”) | 1 | 1 | 9 | 16 |
|  | PS5 (“*Enterobius*”) | 3 | 3 | 4 | 8 |
|  | PS6 (“*Panagrellus*”) | 1 | 1 | 5 | 1 |
|  | Total | 36 | 36 | 73 | 133 |
| Cestodes | *Hymenolepis diminuta* | 7 | 7 | 30 | 55 |
|  | *Hymenolepis nana* | 10 | 10 | 22 | 39 |
|  | Total | 17 | 17 | 51 | 93 |
| Eimeria sp. |  | 15 | 15 | 30 | 54 |
| Ectoparasites | Lice | 45 | 45 | 25 | 46 |
|  | Ticks | 6 | 6 | 0.5 | 1 |
|  | Total | 47 | 47 | 25 | 46 |

Neither microbiota alpha diversity nor richness was related to host variables or most of the parasite presences. The only significant variable was *Eimeria* presence for both diversity (with significance of *p* = 0.038 and the coefficient: 7.8) and richness (*p* = 0.011, coef.: 24.5) (Figure 1). For beta diversity, *H. diminuta* and ectoparasites presence both had significant effects on two of the four metrics (Appendix 2).

**Figure 1:**
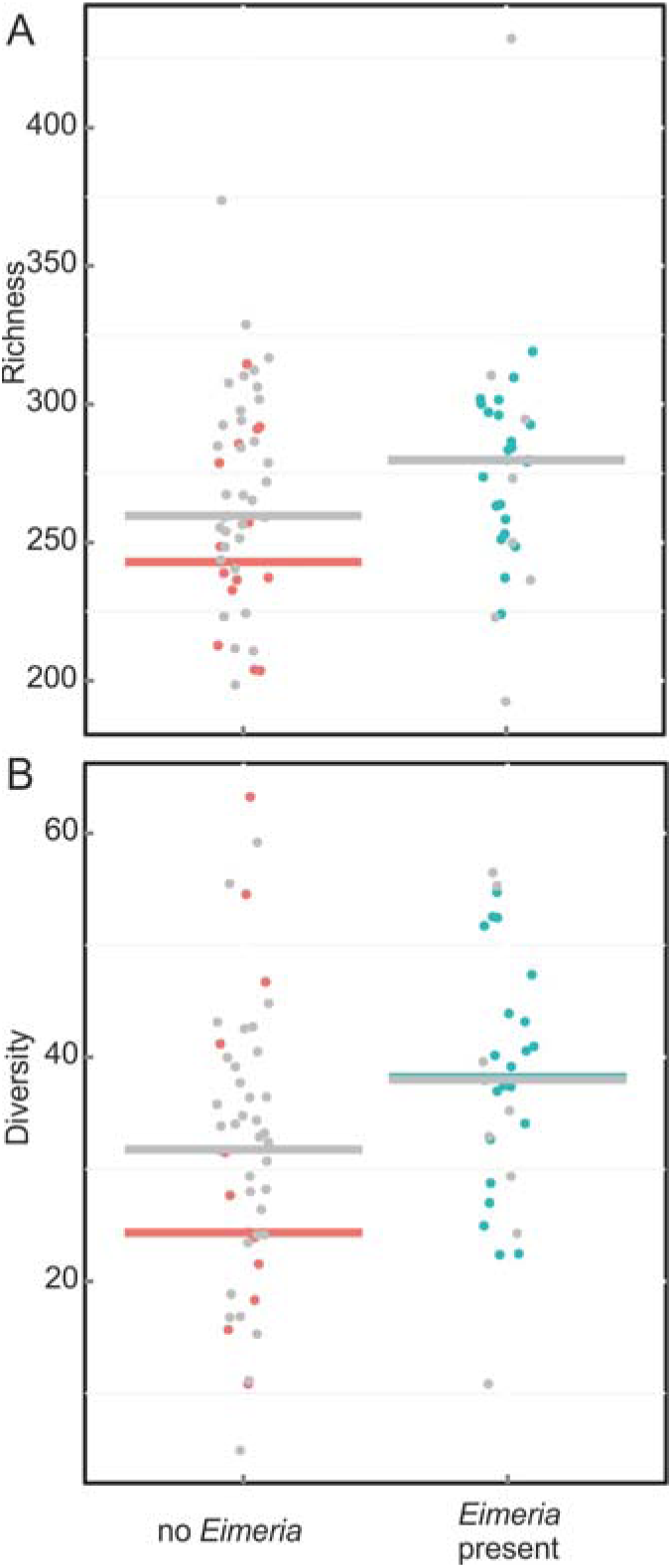
a) Richness and b) inverse Simpson diversity of microbiota in mouse lemurs with and without Eimeria infection. Red and blue colors represent individuals (n=11) which had both negative and positive detections of Eimeria infection, while grey points represent other individuals. The horizontal lines are medians for multiple-sampled individuals and for all individuals.

All the parameter estimates, including associations between species, presented in the following chapters as ‘significant’ have statistical support based on the 90% central credible interval, unless otherwise stated. A positive association between two species means that they occur together more often than expected based on their (dis)similar habitat preferences and purely by random, whereas negative associations implies that they occur together less often than expected based on their habitat preferences or by random.

### Model i) and ii): responses to the environment and associations between parasites species

In model i), all significant associations between parasite species at the level of individual lemurs were positive (Figure 2a). Cestode *Hymenolepis diminuta* had strong *(R* > 0.79, Appendix 4, Figure A2a) positive associations between putative nematodes species 1 and 2, which both had in turn particularly strong positive association with putative nematode species 4. *Eimeria* and *H. diminuta* had a strong association *(R =* 0.84, Appendix 4, Figure A2b) at the level of transects. At the temporal level (Figure 2b), there were both negative and positive associations, meaning that some parasites were co-occurring during the same year (positive) or occurring during different years (negative). These associations coincide with differences in parasite prevalence (Table 1): cestodes were less prevalent in 2011 whereas the prevalence of ectoparasites was more similar between years, with a high prevalence of lice and low of ticks.

**Figure 2:**
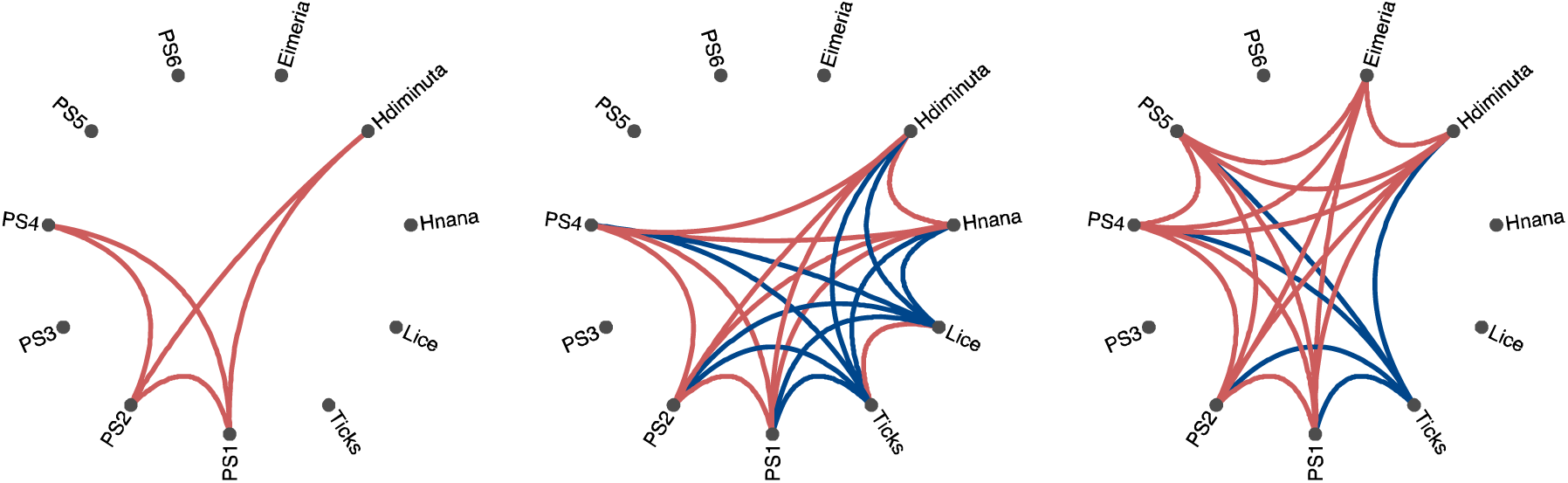
Associations at the level of a) individual lemurs and b) years between different parasite species for the constrained model; and c) associations at the level of individual samples for the unconstrained model. Red lines denote positive associations and blue lines negative associations, all with significant statistical support based on the 90% central credible interval.

There were a few significant explanatory variables for presences of parasite species. *Eimeria* was more probable to be present when the host lemur had better body condition (Appendix 4 Table A1). Lice and ticks were more probable to occur in males, while the occurrence probability of lice was negatively correlated with testis size. Both PS1 and PS4 were negatively correlated with higher age, whereas PS1 was also positively correlated with body condition. Neither the mode of parasite infection – indirect or direct – nor ecto- or endoparasitism significantly explained the differences in responses to parasite species.

In the unconstrained model ii), both the amount of significant associations and the amount of interactive species was the same as with the constrained model at the level of sampling units (Figure 2c) and years (Figure 2b). *Eimeria* and PS5 showed unconstrained association patterns with several nematode species and ticks, but these did not exist after accounting for their habitat requirements (Figure 2a-b). No associations changed directions: positive associations were positive in both models at all levels, as were the negative associations. All the significant constrained associations at the level of individual lemurs (Figure 2a) were also visible in the unconstrained model (Figure 2c), implying that some of the unconstrained associations were due to (dis)similar habitat requirements.

After partitioning the variation explained by the model i), the covariates related to the lemurs accounted for 49% of the total variation explained by the model, whereas the covariates related to seasonality accounted for 9.5% (Appendix 4, Fig. A3a). Species traits explained 46% of the total variation captured with fixed effects (which was 58.8% of the total variation explained by the model). Random effects accounted for 18% at the scale of lemurs, 9.8% at the scale of transects and 13% at the scale of years of the total variance explained by the model.

No traits had significant effects, but there was a strong phylogenetic signal in the species responses to their environment (0.92, see Appendix 3 for details).

### Model iii) and iv): responses to the environment and associations between parasites and microbiota

There were three parasite species with significant associations with bacterial families at the individual mouse lemur level: *Eimeria* sp., *Hymenolepis diminuta* and *H. nana* (Figure 3). Whilst *Eimeria* and *H. diminuta* had a positive association (*R =* 0.79; Appendix 4 Figure A2b), *H. nana* and *H. diminuta* had a negative association (*R* = — 0.89). Consequently, *H. diminuta* and *Eimeria* sp. had mostly negative associations with bacterial families (*R* = — 0.85 — –0.98 and *R =* >—0.78 — –0.79, while the associations of *H. nana* positive (*R =* >0.83 —0.90). All bacterial families, which had positive or negative association with parasite species, had positive associations with each other (ft = 0.79— 0.94). *Eimeria* had in addition two negative associations, with Lactobacillales and Pasteurellales. Only Enterobacteriales had negative association with *H. nana*, while it did not have significant positive association with *H. diminuta.* No explanatory variables were significant, except that of Anaeroplasmatales, which were more abundant in males, nor were there associations at the level of transects.

**Figure 3:**
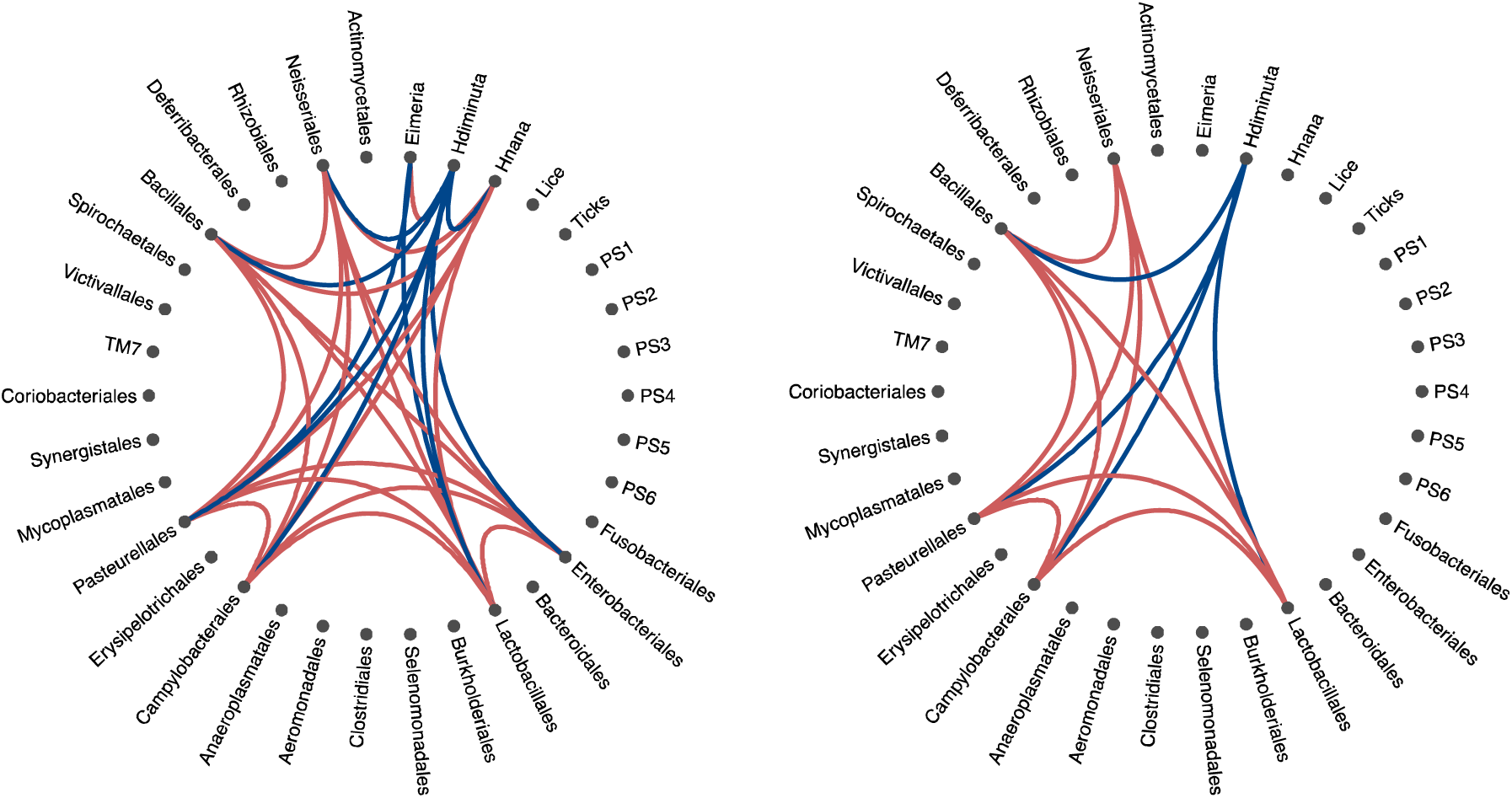
Associations at the level of a) individual lemurs for the constrained model and b) at the level of individual samples for the unconstrained model. Red lines denote positive associations and blue lines negative associations, all with significant statistical support based on the 90% central credible interval.

All the unconstrained associations (Figure 3b) were also visible with the constrained model (Figure 3a). There were fewer significant associations when the environmental covariates were not included in the model, meaning that some associations were not observable before removing the effect of the (dis)similar habitat requirements. Of all the parasites, only *H. diminuta* expressed unconstrained associations (Figure 3b). The negative associations between *H. diminuta* and several bacterial families were present regardless of the inclusion of the environmental constrains, but all the associations between *H. nana* and the bacterial families as well as its negative association with *H. diminuta* were not visible in the unconstrained associations. The species that exhibited any associations with the unconstrained model did so also with the constrained model, with the exception of *Eimeria* and *H. nana*. No associations changed directions between unconstrained and constrained models.

The covariates related to the lemurs accounted for 56% of the total variation explained, whereas the covariates related to seasonality accounted for 15% (Appendix 4 Fig. A3). Species traits explained 44% of the total variation explained with fixed effects (which was 71% of the total variation explained with the model). Random effects accounted for 19% at the scale of lemurs, and 11% at the scale of transects of the variance explained by the model.

No traits had significant effects, but there was a strong phylogenetic signal in the species responses to their environment (with posterior mean 0.94; for more details, see Appendix 3).

## Discussion

Our results show that some intestinal macroparasite species were associated more often than predicted by chance with other helminths, and that they are also associated with differences in microbiota composition. The presence of *Hymenolepis diminuta* – but not the closely related species *Hymenolepis nana* – is correlated with markedly different parasite and microbiota community (Figures 2, 3;. permutational manova: Appendix 2). In more detail, *H. diminuta* presence is negatively associated with bacterial orders Enterobacteriales, Lactobacillales, Campylobacteriales, Pasteurellales, Bacilliales and Neisseriales (Figure 3), whilst it has positive association with nematode species, putatively *Strongyloides* and *Caenorhabditis* (Figure 2a). In addition, *Eimeria sp.* was positively associated with *H. diminuta* and thus also negatively associated with Pasteurellales and Lactobacillales (Figure 3). Surprisingly, host variables did not have significant effects on microbiota, while there were some significant variables affecting the parasite presence.

Two previous studies have looked specifically for the effect of *Hymenolepis* sp. infection: a laboratory study on rats and *H. diminuta* showed a reduction in Lactobacillales and Bacillales and increase in Bacteroidales and Clostridiales, while not showing differences in alpha and beta diversity (McKenney *et al.* 2015), and observational study on wild mice showed both increase and decrease OTUs belonging to Bacteroidales and Clostridiales (Kreisinger *et al.* 2015). Thus, our study is partly consistent with previously found results. Nevertheless, collating OTU data on order level can mask the changes in lower levels: if there has been actually both decrease and increase in different OTUs within an order, say Bacteroidales or Clostridiales, these might not show in upper taxonomic level.

The associations are not necessarily a sign of direct interactions (such as competition) between species, but they can also be driven by indirect causation, like host immune response (host type 2 immunity) towards parasites driving the changes in microbiota or microbiota immunomodulation affecting colonization success of parasites (Ramanan *et al.* 2016). *Eimeria*, unlike other surveyed parasites, is an single-celled intracellular parasite, and the immune reaction normally is type 1 -biased (Cornelissen *et al.* 2009). Thus, it is possible, that *Eimeria* affects microbiota by direct competition, while other have indirect effects. *Eimeria* was the only parasite, which affected the alpha diversity of microbiota (Figure 1). In poultry *Eimeria* reduces the alpha diversity (Stanley *et al.* 2014; Wu *et al.* 2014), while it is less often studied in mammals. Evidence so far seems to indicate that they can either increase or decrease alpha diversity (Bär *et al.* 2015; Ras *et al.* 2015).

Our previous study has shown that there is pervasive within-individual variation in mouse lemur microbiota (Aivelo *et al.* 2016). Thus, it is understandable why we did not identify statistically significant host variables in microbiota variation (Appendix 4 Table A2). Our larger dataset, model i), on parasite occurrence did find host traits which affect parasite presence: better body condition seemed to lead to more probable infection with *Eimeria* and PS1 (putative *Strongyloides*) (Appendix 4 Table A1). This is in line with previous studies with mouse lemurs (Rafalinirina *et al.* 2007). Males more often had ectoparasites, which have been also previously noted (Durden, Zohdy & Laakkonen 2010; Zohdy *et al.* 2012), likely due to more common social interaction between males. Lice prevalence decreased with higher testis volume, which was a surprise, as higher testis volume correlates with higher testosterone levels which in turn can be immunocompromising (Zohdy 2012). Age seemed to lead to lower abundance for two nematode species, which might be caused for example by immunity acquisition (Turner & Getz 2010) or more infected individuals dying younger (Hayward 2013).

The associations between parasites within host individuals were positive in model i) (Figure 2a). *Hymenolepis diminuta* again had associations, though this time positive, with other parasite species, whereas *H. nana* did not have significant associations. This analysis did not find a negative association between *H. diminuta* and *H. nana*, nor a positive association between *H. diminuta* and *Eimeria* sp., due to having larger data set than in combined parasite and microbiota analysis (model iii). The year-level associations were positive between endoparasites on the one hand and ectoparasites on the other hand, while the associations between these groups were negative (Figure 2b). This indicates that endoparasites and ectoparasites have differing dynamics, i.e., when ectoparasites are more common, the endoparasites are less common and other way round. This means that some other factors not captured by our variables, can modulate the parasite prevalence. These could be phenology of insects (Atsalis 2008), which are intermediate hosts for endoparasites, and ambient temperature, which affects sleeping patterns and thus nesting site sharing for mouse lemurs (Schmid & Ganzhorn 2009; Karanewsky & Wright 2015).

As we compare the constrained and unconstrained models, some of the associations are *a)* consistent throughout the levels of observation (such as the positive associations between *Hymenolepis diminuta* and some nematode genera exhibited at all levels of observation with the parasite data set (Figure 2a-c). This implies that these are strong associations not related to the (dis)similar habitat preferences of the species, as they are captured by the model regardless of whether these preferences are accounted for or not; or that there are some unmeasured habitat covariates, to which the species respond (dis)similarly, causing the pattern. Other associations are captured only with *b)* the unconstrained model (such as some of the associations between parasites [Figure 2c]), or *ic)* the constrained model (such as many of the year-level association of parasites [Figure 2b] as well as many of the associations between parasite species and bacterial families [Figure 2a]). The latter case *(c)* implies associations hidden by the (dis)similar responses to habitat, but revealed after these are accounted for, and the former *(b)* suggests associations that are due to these responses to habitat, and disappear after they are accounted for. The associations patterns captured in the manner of *a)* and *c)* can be considered as hypotheses for species interactions, as they are non-random co-occurrence patterns even after accounting for the habitat requirements of the species. Parasite-microbiota studies are further complicated by needing to take into account interactions with the host (Kreisinger *et al.* 2015; Loke & Lim 2015). With the framework used in this study, it is not possible to account for the effects that the parasites and/or microbiome most certainly has on the host directly, but by using host characteristics as explanatory variables, we are controlling for some of the effects of the host on the parasites and microbiota.

Parasite and microbiota traits explained almost half of the variation among the species niches (environmental covariates; Appendix 4 Figure A3). This gives support for idea of niche conservatism (Mouillot *et al.* 2006), as the phylogenetic signal in the species responses to their environment was strong. Moreover, the phylogenetic relationships between the species and the covariance structure of the latent variables at the level of lemurs correlated with each other (Appendix 3), suggesting that there are some similar patterns in the phylogenetic relationships and association between species, and this could be due to some unmeasured, phylogenetically conservative traits, which determine the species niches (in and on the lemurs).

General problems with fecal sampling also need to be taken into account while interpreting the results. For microbiota composition, the composition of fecal matter differ in different parts of the intestinal mucosa (Eckburg *et al.* 2005; Walk *et al.* 2010) and thus fecal sampling could give only a partial picture of what is happening in the gastrointestinal tract. Nevertheless, the helminth effect on microbiota is not confined to helminth niche as such (Kreisinger *et al.* 2015). Parasite prevalence cannot be definitely assessed by nonterminal sampling, as there could be parasites in the intestine, but they might not be laying eggs for a number of reasons (Gillespie 2006; Jorge *et al.* 2013). Our observational sampling also limits how strong statements we can give about causations within the system. Our model is further limited by the low prevalence of many of the parasites, including ticks and putative nematode species 3-6 (Table 1). This influences the estimation of the associations, which is sensitive to the rarity of focal taxa, whereas the estimation of the fixed effects is more robust, as information is being borrowed from other taxa (Ovaskainen & Soininen 2011; Abrego *et al.* 2016b).

In conclusion, we found associations, which can be considered as hypothesis for interactions between intestinal parasites and gut microbiota in observational data set on wild-living mammal. While the presence of unicellular *Eimeria* was linked to a higher alpha diversity, the betadiversity was modulated by ectoparasite and cestode *Hymenolepis diminuta* presence. We also explored the use of novel joints species distribution modelling for identifying interactions between multiple parasite species and microbiota, and found differing responses on microbiota orders by closely related two cestode species and *Eimeria* sp. This investigation should be followed by experimental work in order to establish the causative reasons for variation in the microbiota.

## Author contributions

T.A. conceived and designed the study and collected and processed the samples, A.N. conducted the joint species distribution modelling and both authors analysed and interpreted the data and wrote the article.

## Acknowledgments

We thank Otso Ovaskainen, Juha Laakkonen, Jukka Jernvall and Aura Raulo for helpful discussions, and Kate Ihle on the comments on the manuscript. We also thank Jani Anttila and Alan Medlar for help with bioinformatical and statistical analysis, Andry Herman Rafalinirina and Victor Rasendry for the assistance in the field and Agnés Viherä and Lars Paulin with the sequencing. T.A. was funded by Oskar Öflunds Stiftelse, and A.N. was funded by the Research Foundation of the University of Helsinki.

## Data accessibility

All metadata used has been uploaded to Figshare: doi: 10.6084/m9.figshare.3385573 (Aivelo & Norberg 2016). These are also described in detail in Appendix 1. All sequence data has been deposited to SRA (accession numbers: SRP042187 and SRP063971) and the correspondence to samples can be found in metadata in Figshare. The code for beta diversity analysis is uploaded to GitHub: https://github.com/aivelo/lemur-community and the code for species modelling can be acquired from authors.

